# Sonogenetic control of cardiomyocytes and cardiac pacing using exogenous Transient Receptor Potential A1 channels

**DOI:** 10.64898/2026.01.24.701376

**Authors:** Lei Zhang, Jing Zhang, Tsui-Min Wang, Callum Walsh, Farah Sheikh, Sreekanth H. Chalasani, James Friend

## Abstract

Electronic cardiac pacemakers are the standard of care for treating arrhythmias but, even for wireless pacemakers, they require an intracardiac implant procedure and are invasive. Lead-related complications, including life-threatening infections, and limited suitability in pediatric patients restrict pacemaker use. Biological pacing has emerged as a hardware-free alternative, leveraging cell reprogramming approaches to generate pacemaker-like cardiomyocytes. Despite progress, biological strategies face challenges related to complex signaling pathways, intercellular coupling, durability of effect, efficacy, and safety. In addition, once ion channels or cells are implanted, biological pacemakers offer limited real-time programmability, whereas electronic pacemakers enable continuous adjustment of pacing output based on cardiac performance.

Here we propose a hybrid approach that combines genetic sensitization with external control by exploring the feasibility of sonogenetic cardiac pacing. We express the ultrasound-sensitive ion channel hsTRPA1 in cardiomyocytes and use noninvasive ultrasound to modulate channel activity and cardiac function. Using calcium imaging, we show that hsTRPA1 potentiates ultrasound-evoked responses in cardiomyocytes. Furthermore, cardiac expression of hsTRPA1 in mice increases heart rate in response to ultrasound delivered noninvasively through the intact chest, without evidence of a thermal mechanism under the conditions tested. Together, these results establish a proof of concept for sonogenetic cardiac pacing and define an initial acoustic and genetic parameter space compatible with preserving baseline cardiac function. More broadly, this work positions sonogenetics as a minimally invasive, wireless strategy for cardiac rhythm control and motivates future optimization and safety studies toward translation.

## MAIN

The heart rhythm is coordinated by the cardiac conduction system, in which the sinoatrial (sinus) node initiates electrical impulses that propagate through the atria and ventricles to trigger contraction. Aging, myocardial disease, and congenital or acquired conduction defects can impair impulse generation or propagation, leading to bradyarrhythmias, syncope, reduced cardiac output, and heart failure. For these conditions, implantable electronic pacemakers are the standard of care and have been used clinically since 1958.^1^ Contemporary systems typically require an implanted pulse generator and intracardiac leads placed via transvenous access, with generator replacement every ∽ 5–15 years. Leadless (wireless) pacemakers reduce lead-related failure modes and some infectious complications, but still require intracardiac implantation and do not address all pacing indications.^2^

Despite their effectiveness, electronic pacing remains limited by hardware-associated complications, including device- and lead-related infections,^3^ and by special considerations in patients with congenital heart disease and pediatric populations, who may face repeated interventions over a lifetime.^4,5^ These constraints have motivated the development of biological pacemakers, which aim to restore or replace sinoatrial node function without permanently implanted hardware.^6^ Prior work has pursued four main strategies:^1^ (i) functional re-engineering by viral overexpression of pacemaker-associated ion channels in cardiomyocytes;^7,8^ (ii) cell-based pacing using embryonic stem cell- or induced pluripotent stem cell-derived pacemaker-like cells;^9,10^ (iii) hybrid approaches in which implanted cells deliver ion channels or pacing currents;^11,12^ and (iv) somatic reprogramming, for example via TBX18 expression to induce sinoatrial-like phenotypes in ventricular cardiomyocytes.^13,14^ Although promising in preclinical models, these approaches face challenges in durability, safety, and controllability, and generally offer limited real-time programmability after implantation.

Here we explore a hybrid strategy that couples genetic sensitization with external, noninvasive control. Sonogenetics, the use of ultrasound to modulate genetically specified ion channels,offers a potential route to programmable, wireless control of cardiac excitability without chronically implanted electrodes.^15^ Building on this concept, we test whether cardiomyocyte expression of the ultrasound-sensitive channel hsTRPA1 can enable on-demand modulation of cardiac activity using ultrasound delivered through intact tissue.

Sonogenetics uses ultrasound to noninvasively modulate cellular activity through genetically specified actuators, most commonly ion channels that couple acoustic energy to cation influx and downstream excitability.^16^ The dominant transduction mechanisms are typically described as both thermal and mechanical: thermosensitive channels can be gated by ultrasound-induced heating, whereas mechanosensitive channels respond to ultrasound-driven stresses and deformations at the cell membrane.^17^ Proposed mechanical models include intraleaflet cavitation^18^ and direct membrane deflection; consistent with the latter, high-speed digital holographic microscopy has captured real-time membrane motion during ultrasound exposure sufficient to generate charge effects observed in experiments.^19^

The concept was introduced in 2015, when Ibsen and colleagues reported ultrasound-evoked behavioural responses in *C. elegans* upon overexpression of the mechanosensitive TRP– 4 channel.^20^ Subsequent work has extended sonogenetic control toward mammalian systems by identifying and engineering actuators that couple ultrasound to rapid ion flux and cellular activation. Examples include ultrasound-driven gene regulation using heat-shock promoters to control chimeric antigen receptor expression in T cells^21^ and sonogenetic neuromodulation using engineered mechanosensitive channels to modulate subthalamic circuitry in Parkinsonian mouse models.^22^

Recent studies also support the feasibility of ultrasound-based cardiac pacing. For example, Marquet and colleagues demonstrated pacing in a porcine model using high-intensity focused ultrasound, reporting ultrasound-evoked premature ventricular depolarizations detected as premature QRS complexes via electrocardiography.^23^ In these and related approaches, pacing is achieved without genetically encoded ultrasound actuators; consequently, relatively high acoustic intensities are often required to achieve reliable excitation in vivo, raising concerns about off-target bioeffects and tissue injury. To reduce excitation thresholds, some studies administer ultrasound contrast agents or other sensitizers, which can enhance mechanical effects^24^ but introduce additional procedural complexity and potential safety constraints. Sonogenetics offers a complementary strategy in which expression of an ultrasound-sensitive ion channel lowers the acoustic threshold required for excitation, thereby expanding the safety margin for in vivo stimulation. In addition, restricting actuator expression to defined cardiac cell populations can improve spatial specificity beyond what is achievable with ultrasound focusing alone, enabling cell-type-selective modulation of cardiac excitability.

A suitable genetic toolkit is essential for applying sonogenetics to cardiac pacing, because the choice of actuator determines the dominant transduction mechanism, stimulation thresh-old, and potential off-target effects. Several classes of ultrasound-responsive actuators have been reported. Mechanosensitive TRP channels provided an early demonstration that ultrasound can drive neuronal activation and behavioral outputs.^20^ Bacterial mechanosensitive channels such as MscL have been used to enable ultrasound-evoked calcium entry and action potentials in neurons.^25^ Thermosensitive TRPV1 supports a “sonothermogenetics” strategy that leverages ultrasound-induced heating.^26^

Here we focus on transient receptor potential ankyrin 1 (TRPA1). Recent studies report that TRPA1 exhibits comparatively high ultrasound sensitivity among screened candidate actuators, and that the human ortholog (hsTRPA1) is among the most responsive variants.^27^ TRPA1 is a calcium-permeable, non-selective cation channel in the TRP superfamily, enriched in subsets of nociceptive primary sensory neurons and originally identified as ANKTM1 in pain-related pathways.^28^ The channel can be activated by diverse chemical and physical stimuli, including electrophilic irritants, cold, and mechanical forces,^29,30^ leading to membrane depolarization and calcium influx^31^. Although TRPA1 is endogenously enriched in subsets of nociceptive sensory neurons, heterologous expression can repurpose its calcium-permeable, non-selective cation conductance as an actuator in other excitable cells, motivating its evaluation as a means to confer ultrasound responsiveness to cardiomyocytes.

Prior work also suggests that TRPA1 activation can modulate cardiomyocyte excitation– contraction coupling. Andrei and colleagues reported increased calcium influx and altered sarcomere dynamics in isolated mouse ventricular cardiomyocytes following pharmacological activation of TRPA1 with allyl isothiocyanate (AITC).^32^ Motivated by these observations, we hypothesized that sonogenetic activation of hsTRPA1 could provide a noninvasive route to increase cardiac excitability and heart rate, establishing a potential foundation for a programmable, wireless pacing strategy.

## RESULTS

### hsTRPA1 functional expression in cardiomyocytes

To test whether hsTRPA1 can be functionally expressed in adult mouse ventricular cardiomyocytes *in vivo*, we used an adeno-associated virus (AAV) strategy to drive cardiac expression of hsTRPA1. We packaged Myc-tagged hsTRPA1 (AAV9-*cTnT::hsTRPA1-Myc*) in the cardiotropic AAV9 serotype under the cardiac troponin T (cTnT) promoter. Wild-type mice received intraperitoneal AAV injection on postnatal day 3 at a dose of 6 *×*10^13^ genome copies per kilogram and were analyzed at 4 weeks of age (Fig. 1a). We verified hsTRPA1 expression in cardiac tissue by reverse transcription quantitative polymerase chain reaction (RT–qPCR) and by immunofluorescence staining for the Myc epitope in cardiomyocytes. The viral dose and neonatal delivery paradigm were selected based on prior work using neonatal AAV administration in a mouse model of catecholaminergic polymorphic ventricular tachycardia.^33^ Unless otherwise noted, all subsequent emphin vivo experiments used the same neonatal AAV delivery strategy and were performed four weeks after injection.

**FIG. 1.**
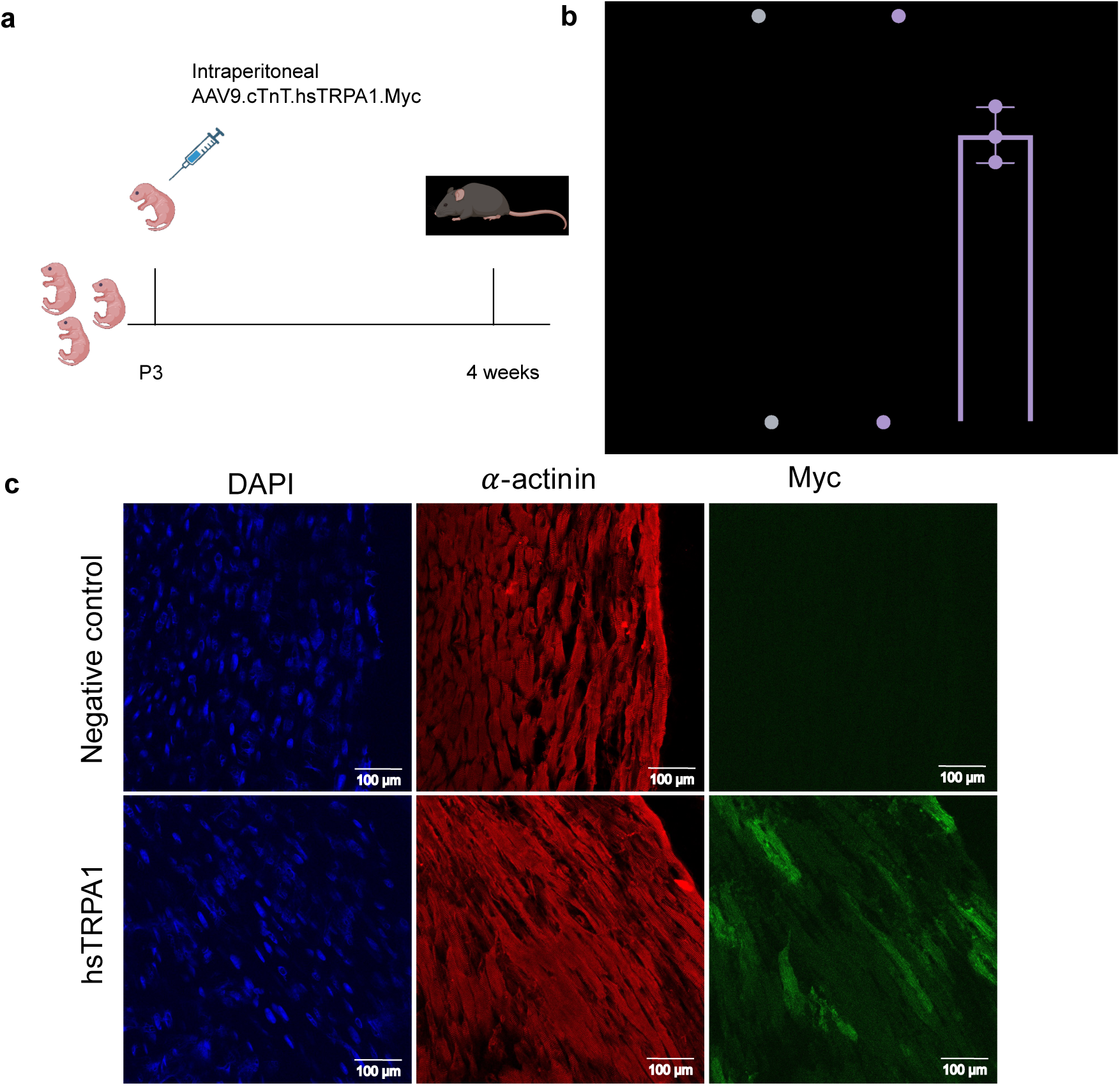
a, Early neonatal AAV delivery strategy and analysis time point four weeks later. b, RT-qPCR analysis of hsTRPA1 and mmTRPA1 levels in wild-type mice and AAV9-hsTRPA1 injected wild-type mice. Results are normalized to GAPDH. Data are presented as mean*±*s.e.m. One-way ANOVA with Tukey’s multiple comparison test. *\*P <* 0.05 (*P =* 0.003 for mmTRPA1 in wild-type mice versus hsTRPA1 in AAV9-hsTRPA1 mice; *P =* 0.002 for mmTRPA1 versus hsTRPA1 in AAV9-hsTRPA1 mice). c, Representative immunofluorescence staining of DAPI (blue), *α*-actin (red) and Myc (green) at four weeks post injection in both uninfected wild-type control mice and wild-type mice injected with AAV9-hsTRPA1. Experiments were repeated independently three times with similar results.

Immunofluorescence analysis revealed robust Myc signal within *α*-actinin^+^ cardiomyocytes in AAV-hsTRPA1-treated hearts, whereas uninjected controls showed no specific Myc staining (Fig. 1c), consistent with cardiomyocyte-restricted expression following AAV delivery. Reverse transcription quantitative PCR (RT–qPCR) further supported these observations, demonstrating a significant increase in hsTRPA1 transcript levels in AAV9-hsTRPA1-injected mice relative to controls (Fig. 1b). RT–qPCR also indicated that endogenous mouse *Trpa1* (mmTRPA1) expression in whole-heart tissue was at or near background levels under our assay conditions. This contrasts with prior reports of endogenous TRPA1 expression in murine cardiomyocytes.^34–36^ Because our experiments primarily compare hsTRPA1-expressing hearts to matched controls processed in parallel, any low-level endogenous mmTRPA1 expression would be expected to affect baseline responsiveness in controls but would not confound interpretation of hsTRPA1-dependent effects. We return to this point when interpreting control responses in subsequent experiments.

To test whether virally delivered hsTRPA1 is functionally expressed in cardiomyocytes, we challenged isolated adult ventricular cardiomyocytes with the TRPA1 agonist allyl isothiocyanate (AITC) and measured intracellular calcium responses. Cells were loaded with the calcium indicator Fluo-4 and fluorescence was recorded continuously. After establishing a baseline, 2 *µ*L AITC (100 *µ*M) was applied during imaging directly at the observed cells by using a pipette (Fig. 2a). Cardiomyocytes isolated from AAV9-hsTRPA1 mice exhibited larger AITC-evoked increases in Fluo-4 fluorescence than cardiomyocytes from wild-type controls (Fig. 2b) 31 cells were observed from 4 mice for both WT and hsTRPA1 groups separately. Peak fluorescence after AITC application was significantly higher in hsTRPA1-expressing cells, whereas most wild-type cardiomyocytes showed no detectable response (Fig. 2c), consistent with functional hsTRPA1 expression. Not all cardiomyocytes isolated from AAV9-hsTRPA1-treated mice responded to AITC, which is consistent with incomplete transduction and/or heterogeneous transgene expression; thus, non-responding cells in this group cannot be assumed to express hsTRPA1.

**FIG. 2.**
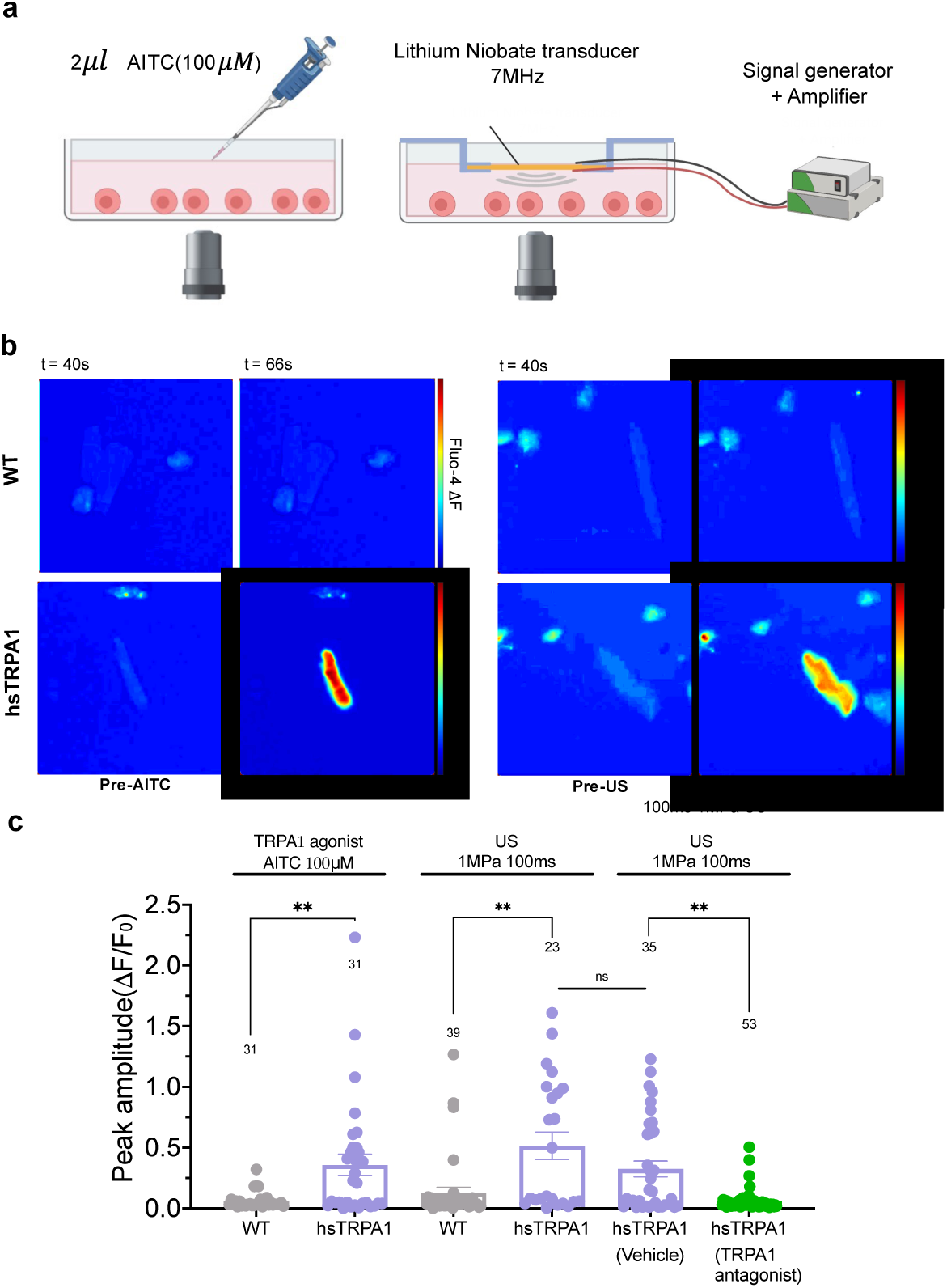
a. In vitro delivery of AITC and the use of a 7 MHz lithium niobate transducer for ultrasound stimuli of the cells. b. Fluo-4 fluorescence signal in isolated adult ventricular cardiomyocytes from wildtype mice and AAV9-hsTRPA1 injected mice before and after treatment with AITC (100 *µ*M) or 100 ms ultrasound pulse stimulation. c. Fluo-4 peak amplitude in isolated adult ventricular cardiomyocytes from wild-type mice and AAV9-hsTRPA1 mice stimulated with TRPA1 agonist (AITC, 100 *µ*M), ultrasound alone, vehicle control (0.1% DMSO) or TRPA1 antagonist (HC–030031, 40 *µ*M). Data are mean *±* SEM; *n =* 4 biologically independent animals. *\*\*P <* 0.01 by two-tailed Mann-Whitney test. The number of cells analyzed is indicated above each bar. Source data are provided with this paper.

### hsTRPA1 potentiates ultrasound responses in cardiomyocytes

Next, we tested whether ultrasound stimulation can evoke hsTRPA1-dependent calcium responses in cardiomyocytes. We built a custom lithium niobate transducer (resonance frequency, 6.776 MHz) for use in the calcium-imaging configuration. Lithium niobate exhibits no hysteresis and can therefore reduce transducer self-heating during electrical-to-mechanical energy conversion.^37^ Acoustic pressure output and associated temperature changes at the sample plane were characterized using a fiber-optic hydrophone in the imaging set-up (Fig. S1a,b). Unless otherwise noted, *in vitro* ultrasound stimulation consisted of only one 100 ms pulse at a peak negative pressure of 1 MPa, which produced an estimated temperature increase of *∼* 0.15 *^°^*C under our conditions (Fig. S1b). The transducer was positioned above the cells during imaging (Fig. 2a).

Ultrasound elicited larger calcium transients in cardiomyocytes isolated from AAV9-hsTRPA1 mice than in cardiomyocytes from wild-type controls (Fig. 2b). Consistent with an actuator-dependent effect, ultrasound-responsive cells were observed more frequently in the hsTRPA1 group, whereas most wild-type cells did not respond detectably to the same stimulus (Fig. 2c). Because AAV transduction was not uniform across all isolated cardiomyocytes, the observed response fraction in the hsTRPA1 group likely underestimates the fraction among hsTRPA1-expressing cells.

To test specificity, we pharmacologically inhibited TRPA1 using the antagonist HC-030031.^38^ In hsTRPA1-treated cardiomyocytes, HC-030031 reduced ultrasound-evoked calcium responses relative to vehicle control, supporting the conclusion that ultrasound responses were mediated by hsTRPA1 channel activity. Together, these data indicate that hsTRPA1 expression potentiates ultrasound-evoked calcium influx in isolated adult ventricular cardiomyocytes.

### hsTRPA1 confers ultrasound sensitivity *in vivo*

We next asked whether hsTRPA1 can enable sonogenetic modulation of heart rate *in vivo*. Translation from the *in vitro* imaging configuration to intact animals requires accounting for acoustic attenuation in overlying tissues. In the *in vitro* set-up, ultrasound propagated through a thin column of aqueous medium between the transducer and the cells, for which attenuation is minimal at the distances used. In contrast, *in vivo* stimulation requires transmission through skin, subcutaneous tissue, and the thoracic cage before reaching the heart, resulting in substantially greater attenuation and potential distortion of the acoustic field. Ultrasound attenuation is commonly quantified by an attenuation coefficient *α*, which depends on acoustic frequency *f* and tissue composition.^39^ Accordingly, we measured the acoustic pressure at the heart using a fiber-optic hydrophone and incorporated tissue attenuation *α* as follows when selecting stimulation parameters for *in vivo* experiments:

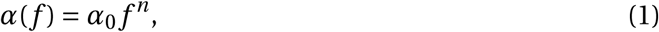

where *α*_0_ is the attenuation constant and *n* is the power-law exponent determined by the tissue’s properties. For *in vivo* electrocardiography (ECG) experiments, we used a lower ultrasound carrier frequency (668 kHz) delivered with a PZT transducer to improve transmission through the thorax. We quantified acoustic pressure output and accompanying temperature changes at the level of the heart using an *ex vivo* measurement configuration with a fiber-optic hydrophone (Fig. S2a–c). Based on these measurements, we selected peak negative pressure amplitudes of 1, 2, and 3.5 MPa for subsequent *in vivo* stimulation. The corresponding temperature increases at the heart were 0, 0.3, and 1.0*^°^*C after 1 min of exposure for 1, 2, and 3.5 MPa, respectively. These temperature increases fall within the ranges reported to be compatible with safe cardiac ultrasound exposure.^40^

Ultrasound was delivered transcutaneously to the hearts of anesthetized mice using a PZT transducer coupled to the chest with ultrasound gel to ensure acoustic transmission (Fig. 3a). ECG was recorded continuously while ultrasound pulses (100 ms every 5 s) were applied for 1 min. Under these conditions, hsTRPA1-expressing mice exhibited a significant increase in heart rate relative to baseline during 3.5 MPa stimulation, whereas wild-type mice showed no significant change (Fig. 3d).

**FIG. 3.**
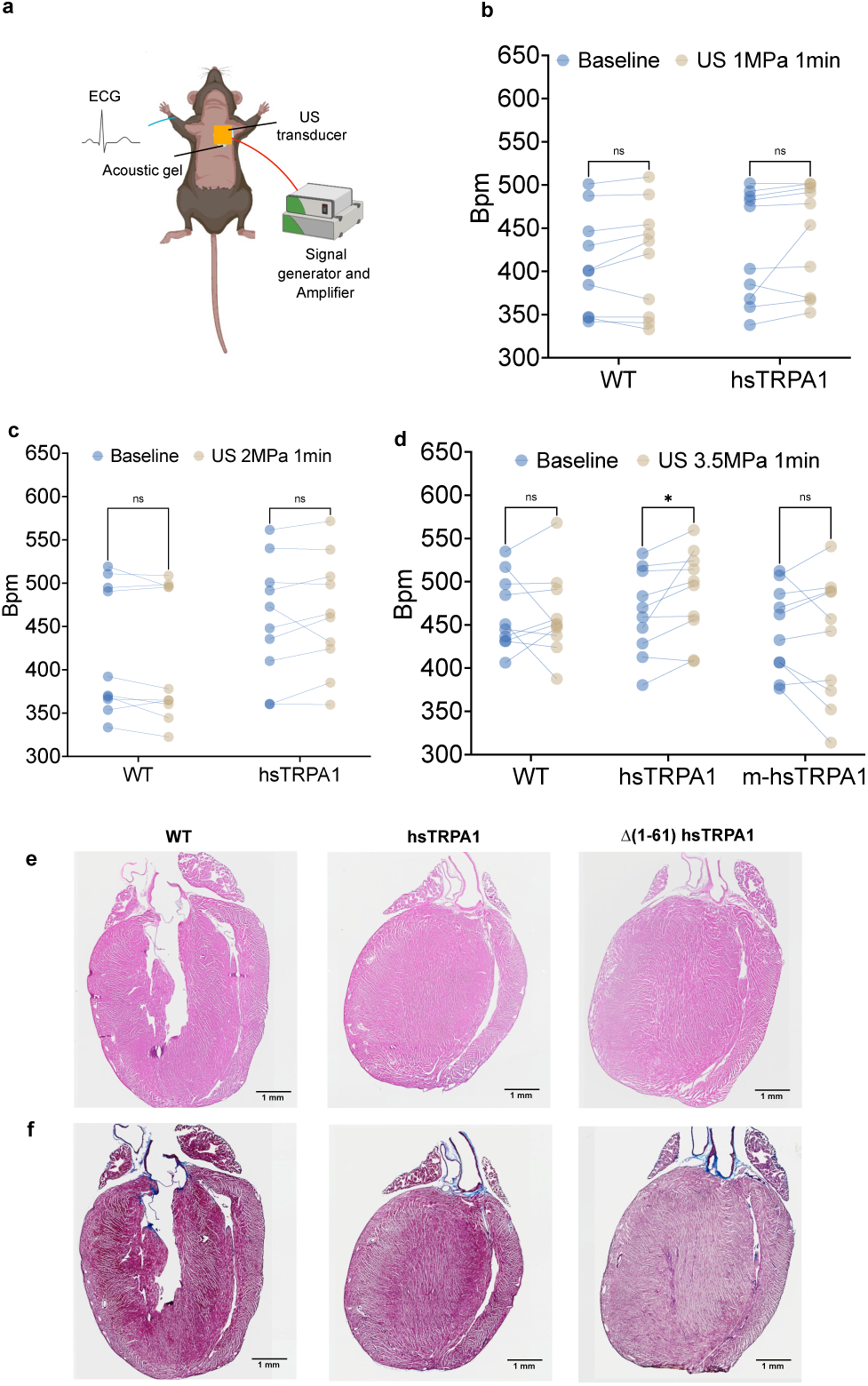
a. Mouse ECG experiment. b,c. Quantification of heart rate in baseline before and after 1 min US stimulation with 1 MPa and 2 MPa for wild-type mice and AAV9-hsTRPA1 mice. d. Quantification of the baseline heart rate before US stimulation, followed by a measurement after 1 min US stimulation for wild-type mice, AAV9-hsTRPA1 mice and mutant AAV9-Δ1-61 hsTRPA1 mice. In these experiments, *n =* 10 (b c d) with biologically independent animals. Data are presented as pairwise as before and after US stimulation for one mouse; *\*P <* 0.05 by Wilcoxon matched-pairs signed rank test with Holm–Šidák correction for multiple comparisons. e,f, Representative cardiac sections stained with hematoxylin and eosin (e) and Masson’s Trichrome staining (f) from WT, hsTRPA1 and Δ(1–61) hsTRPA1 mice at weeks 2 after ultrasound stimulation. Scale bars, 1 mm.

To test whether this effect required hsTRPA1 gating, we evaluated a loss-of-function hsTRPA1 variant lacking the intracellular N-terminal tip domain (Δ(1–61) hsTRPA1). This 61-amino-acid region is highly conserved in mammals and has been reported to be necessary for ultrasound sensitivity; deleting it abolishes ultrasound-evoked responses while preserving other aspects of channel function.^27^ Consistent with an hsTRPA1-dependent mechanism, mice expressing Δ1–61 hsTRPA1 did not exhibit a significant heart-rate increase under the same 3.5 MPa stimulation paradigm (Fig. 3d).

Ultrasound-evoked heart-rate modulation was pressure dependent. At 1 and 2 MPa, stimulation for 1 min did not produce significant heart-rate changes relative to baseline in either wild-type or hsTRPA1-expressing mice (Fig. 3b,c). For 3.5 MPa, the slight higher temperature increase as 1 *^°^*C after 1 min exposure did not affect the heart rate of WT and Δ1–61 hsTRPA1 mice, confirming the heart rate increasing in hsTRPA1 mice group were not caused by heat. Together, these data indicate that non-thermal, hsTRPA1-dependent increases in heart rate are observed within a restricted acoustic regime and underscore the need to control pressure, duty cycle, and exposure duration to limit ultrasound-induced heating in future applications.

To assess whether ultrasound exposure altered cardiac morphology or produced tissue damage, we performed histological analysis of hearts harvested 2 weeks after stimulation. Hematoxylin and eosin staining of whole-heart sections revealed no gross differences in chamber dimensions or myocardial integrity in all wild-type mice, AAV9-hsTRPA1-treated mice and Δ1–61 hsTRPA1 mice (Fig. 3e). To assess collagen remodeling and fibrotic responses, adjacent sections were stained with Masson’s trichrome. Trichrome staining revealed no apparent increase in interstitial, perivascular, or replacement fibrosis(Fig. 3f), providing initial evidence that the stimulation paradigms used here did not produce overt structural injury.

## DISCUSSION

This study establishes hsTRPA1 as a sonogenetic actuator for cardiac modulation, spanning ultrasound-evoked calcium entry in isolated adult ventricular cardiomyocytes *in vitro* and transcutaneous, ultrasound-driven increases in heart rate *in vivo*. Together, these results expand the sonogenetic toolkit toward cardiac applications and support the feasibility of a genetically enabled, externally addressable strategy for rhythm modulation that does not require chronically implanted electrodes.

At the cellular level, hsTRPA1 expression increased the probability and amplitude of ultrasound-evoked calcium transients, and pharmacological antagonism reduced these responses, supporting an hsTRPA1-dependent mechanism. In parallel, control cardiomyocytes exhibited minimal ultrasound responsiveness under the same conditions, and responses in controls were not reduced by TRPA1 blockade, arguing against an endogenous TRPA1-mediated mechanism at the stimulation settings tested. Our RT–qPCR measurements also indicated that endogenous mouse TRPA1 transcript levels in whole-heart samples were at or near background. Prior literature regarding TRPA1 in the myocardium remains mixed: some studies report TRPA1-associated signaling and functional effects in cardiac preparations, whereas others find minimal TRPA1 expression and no biologically relevant functional TRPA1 in primary murine cardiomyocytes, underscoring strong dependence on cell type, preparation, and assay sensitivity.^34,41^ This uncertainty does not undermine the present approach, which relies on controlled hsTRPA1 expression, but it motivates the careful electrophysiological characterization of downstream consequences of hsTRPA1 opening in cardiac tissue, including effects on action potential morphology, refractory behavior, and susceptibility to arrhythmia.

At the level of the organism, hsTRPA1 enabled an increase in heart rate in response to transcutaneous ultrasound delivered through the intact chest, and a loss-of-function hsTRPA1 variant (Δ1–61) that abolishes ultrasound sensitivity prevented this effect, providing genetic evidence that hsTRPA1 gating is required.^27^ Responses were strongly dependent on acoustic settings: lower pressures did not produce measurable heart-rate changes, whereas higher-intensity exposures increased heart rate while maintaining a small amount of heat. This delineates an initial parameter space in which hsTRPA1-dependent effects can be separated from non-specific thermal effects and highlights the need to control peak pressure, duty cycle, and exposure duration when designing cardiac sonogenetic protocols. More broadly, these findings align with the extensive literature on mechano-electric coupling in the heart, in which mechanical perturbations can alter cardiac electrophysiology and rhythm.^42,43^

Ultrasound modulation of cardiac rhythm also has substantial precedent without genetic sensitization. Focused ultrasound has been shown to elicit premature ventricular depolarizations and pacing-like events in large-animal models.^23^ A first-in-human feasibility study further reported that transcutaneous ultrasonic left ventricular pacing, facilitated by an echocardiographic contrast agent, can achieve repeated paced beats and appears safe under monitored conditions.^44^ These studies support ultrasound as a viable physical modality for cardiac stimulation, but they also emphasize important limitations of non-genetic ultrasound pacing, including reliance on microbubble contrast agents and limited molecular or cellular specificity.^23,43^ By contrast, hsTRPA1 introduces specificity through genetically restricted expression, potentially enabling stimulation selectivity that is biologically encoded rather than achieved solely by acoustic focusing.

From a translational perspective, hsTRPA1-based sonogenetic pacing is best viewed as complementary to existing device-based and biological pacing strategies. Leadless electronic pacemakers already reduce lead-associated complications but still require intracardiac implantation and remain constrained by device form factor, battery longevity, and indication-specific limitations.^45^ A sonogenetic approach, if made sufficiently efficient and safe, could offer on-demand, external control without permanently implanted electrodes, and could in principle be paired with cell-type-specific promoters or delivery strategies to target defined cardiac cell populations. Key next steps include (i) defining electrophysiological effects and arrhythmia risk across a wider range of acoustic parameters; (ii) quantifying transduction efficiency and expression stability across time; and (iii) developing closed-loop control strategies that couple ultrasound delivery to real-time physiological sensing. Addressing these points will be essential to evaluate this approach’s feasibility beyond proof-of-concept and to motivate future translational studies.

## METHODS

### Ethical Statement

All research and animal procedures were in full compliance with the ethical guidelines of the Institutional Animal Care and Use Committee of the University of California San Diego, ethical regulations established by the Institutional Biosafety committee of the Salk Institute for Biological Studies.

### Generation of experimental animals

Studies were performed using a total of 54 adult mice, including both males and females. Animals were group housed at 21–23*^°^*C at a humidity of 45 - 55%. The temperature and humidity were monitored and a 12 h–12 h light-dark cycle was maintained, alternating at 6:00 and 18:00. All *invitro* and *invivo* data were collected after four weeks of virus injection.

### Sequences and adeno-associated virus vectors

Human TRPA1 peptide sequences were retrieved from the National Center for Biotechnology Information (NCBI) RefSeq database (Homo sapiens; NCBI Taxonomy 9606; RefSeq XP_016869435.1). Adeno-associated virus vectors expressing either hsTRPA1 or ankyrin TRPA1 mutants were designed and synthesized, containing a cytomegalovirus promoter and Myc tag (Vector Builder). Adenovirus was packaged and grown through the GT3 Core at Salk Institute of Biological Studies as previously described.^27^ The mutant constructs generation of ankyrin TRPA1 was performed as previously described^27^ with some specific modifications. Briefly, a PCR-based approach was used to delete the aa of 1–61 and nucleotide of 1–182.

### Adeno-associated virus injections

Early AAV injections were performed on postnatal day 3 to deliver 6*×*10^13^ genome copies per kilogram in 50 *µ*l of saline solution containing AAV9-hsTRPA1 or AAV9-mutant-hsTRPA1. Virus was delivered by using a 31-gauge needle and syringe (Monoject, 8881600800) via intraperitoneal injection. Successful viral delivery was confirmed post-mortem via immunohistochemistry for the Myc-tag.

### Mouse adult ventricular myocytes isolation

Ventricular myocytes were isolated from adult mouse ventricles were performed as previously described, with some modifications.^46–48^ Briefly, single ventricular myocytes were isolated from mouse ventricles using the enzymatic digestion method by Langendorff. Hearts from heparinized (100 U/mL, intraperitoneal injection) mice were each washed in and perfused with 4 mL of warmed Tyrode’s solution (137 mM NaCl/ 0.5 mM MgCl_2_/ 1.8 mM CaCl_2_/ 4 mM KCl/10 mM HEPES/ 5 mM Glucose; pH 7.4 with NaOH). Each heart was moved to the to the Langendorff system and perfused for 3–5 minutes with calcium-free Tyrode’s (137 mM NaCl/ 0.5 mM MgCl_2_/ 1 mM MgSO_4_/ 1 mM KH_2_PO_4_/ 4 mM KCl/ 10 mM HEPES/ 5 mM Glucose; pH 7.2 with NaOH), followed by a digestion solution (1.5 mg/mL Type II Collagenase and 0.1 mg/mL Protease) for 10–12 min, and a subsequent perfusion with calcium-free Tyrode’s for 3–5 min to flush out the enzymes. After placing the digested heart in a Petri dish containing a 1:1 ratio of Normal Tyrode’s and calcium-free Tyrode’s solutions (8 mL), isolated ventricular myocytes were minced into pieces. Rod-shaped ventricular myocytes were found to settle down and remain alive for up six hours for *in vitro* calcium imaging experiments.

### In vitro pharmacology

For activation of TRPA1, we added 2 *µl* AITC (100 *µ*M in DMSO, allyl isothiocyanate; Sigma-Aldrich # 377430) to the targeted cells. For inhibition of TRPA1, we incubated cells with the antagonist HC–030031 (40 *µ*M in DMSO; Cayman Chemicals #11923) for 45 min before ultrasound simulation. All drugs were diluted with assay buffer in the fluo-4 NW calcium assay kit (ThermoFisher scientific, F36206). The final concentration of DMSO in the solution was 0.1% or lower for all groups. A 0.1% DMSO solution was also used as the vehicle control.

### Calcium imaging system

To measure the in-cell calcium signaling from agonist and US stimulation or antagonist inhibition, the fluo-4 AM (Fluo-4 NW Calcium Assay Kit; ThermoFisher scientific, F36206) was incubated with cells for 30 min at room temperature. The calcium change over time of individual mice ventricular cells were indicated by the fluorescence intensity change, acquired using a fluorescence microscope (ThermoFisher Scientific, EVOS M7000 IMAGING SYSTEM). The captured videos were collected using a 10x objective at 20 frames per second with a GFP filter. Our custom-made lithium niobate transducer with a 6 mm transparent circular window in diameter was immersed in the solution above the cells, which did not affect the light path of the inverted fluorescence microscope. All experiments were performed at a room temperature of approximately 22*^°^*C.

### Calcium imaging analysis

All calcium imaging analyses were performed using custom scripts written in MATLAB. Freshly isolated adult mouse ventricular cardiomyocytes typically settled as rod-shaped, cross-striated cells and were quiescent at baseline. Cells were segmented, and Fluo-4 fluorescence intensity was quantified over time for each cell (defined for use later as a region of interest, ROI) and exported to .csv files. For each frame, the script converted the image to grayscale, computed the mean pixel intensity within each ROI, and then subtracted background fluorescence measured from a cell-free region to correct for nonuniform illumination and video-to-video imaging variability.

Calcium signals were expressed as Δ*F* /*F*, where *F* denotes the background-subtracted ROI fluorescence and *F*_0_ denotes the mean fluorescence during the first 1 min baseline period acquired before drug addition or ultrasound stimulation. Peak Δ*F* /*F* amplitude was calculated for each cell within the 30 s interval following pharmacological treatment or ultrasound stimulation. Unpaired, nonparametric comparisons were performed using a two-tailed Mann– Whitney test, and *P* values were calculated accordingly.

### Immunocytochemistry

Isolated heart samples were washed in KCl (0.3 M) phosphate-buffered saline (PBS) and cryoprotected in 30% (w/v) sucrose in PBS overnight at 4 *^°^*C. Tissues were then cryosectioned at 10 *µ*m thickness onto glass slides and stored at −80*^°^*C. Heart cryosections were fixed in 100% acetone (Fisher Scientific, A18-1) at *−*20*^°^*C for 10 min, then blocked in 5% donkey serum (Sigma-Aldrich, D9663) prior to antibody incubation.

Sections were then incubated overnight at 4 *^°^*C with primary antibodies against sarcomeric *α*-actinin (mouse monoclonal anti-*α*-actinin, Sigma-Aldrich, A7811) and Myc-tagged hsTRPA1 (*Myc-Tag* (71D10) rabbit monoclonal antibody, Cell Signaling Technology #2278). Secondary antibody staining was performed at room temperature for 45 min using donkey anti-mouse IgG (H+L) Alexa Fluor 555 (1:400, Thermo Fisher Scientific, A31570) and donkey anti-rabbit IgG (H+L) Alexa Fluor 488 (1:400, Thermo Fisher Scientific, A21206). Immunofluorescence images were acquired at 20*×*using a Leica SP8 confocal microscope.

### RNA analysis

Total RNA was extracted from heart tissue using TRIzol (Invitrogen) according to the manufacturer’s instructions. RNA concentration was quantified using a NanoDrop spectrophotometer (Thermo Fisher Scientific). First-strand cDNA was synthesized using the iScript cDNA Synthesis Kit (Bio-Rad, #1708891). RT–qPCR was performed using a universal qPCR master mix (Luna, M3003L) with primers obtained from Integrated DNA Technologies. Primer sequences were as follows: forward 5’-CCG CTT ACA GCC CTC AAC G -3’ and reverse 5’-AGC TCT AAA TCC ATA AGC CAA CC -3’ for hsTRPA1 and Δ1–61 hsTRPA1, forward 5’-GAG AGT GTT TCC TCG TCC CG -3’ and reverse 5’-ACT GTG CCG TTG AAT TTG CC -3’ for GAPDH, forward 5’-CAC AGA CCG ACT AGA TGA AGA AG -3’ and reverse 5’-CAG GAG GAT GTC AGG ATT GT -3’ for mmTRPA1. RT–qPCR was performed on heart cDNA using Power SYBR Green PCR Master Mix (Applied Biosystems, 4309155) on a Bio-Rad thermocycler. Relative gene expression was normalized to GAPDH mRNA levels, as indicated. PCR products were confirmed by Sanger sequencing (Eton Biosciences).

### Ultrasound pressure and temperature measurements

Ultrasound pressure and temperature were measured at the same spatial location and using the same experimental geometry as in the corresponding stimulation experiments. Acoustic pressure was recorded using a fiber-optic hydrophone (Precision Acoustics) connected to a digital oscilloscope (Tektronix TBS 1052B) with acquisition controlled by a computer (Lenovo ThinkPad Ultrabook). For *in vitro* calcium imaging experiments, the hydrophone fiber tip was positioned at the bottom of the Petri dish at the plane containing the adult ventricular myocytes, aligned with the acoustic axis of the lithium niobate transducer via a small access hole in the dish bottom. Deionized water was added to reproduce the acoustic coupling and transmission conditions used in the imaging experiments. For *in vivo* ECG experiments, the transducer was coupled to the anterior thoracic skin overlying the heart using ultrasound gel, consistent with the ECG stimulation setup. The dorsal surface was then surgically opened, and the fiber-optic hydrophone was positioned at the anterior surface of the heart to measure acoustic pressure transmitted through the skin and ribs.

### Ultrasound transducer

For *in vitro* calcium imaging experiments, a custom piezoelectric transducer was fabricated from a 10 *×* 10 *×* 0.5 mm single-crystal lithium niobate (LiNbO_3_) substrate, diced from a 4-inch double-side polished wafer (Precision Micro-Optics LLC, Burlington, MA, USA) with a 128*^°^*Y-rotated X-propagation orientation. Electrodes were deposited on both faces of the wafer prior to dicing. Briefly, 6 mm-diameter polyimide tape circles (CS Hyde Company, Lake Villa, IL, USA) were affixed and aligned on the two wafer surfaces to define a transparent aperture. A 20 nm chromium adhesion layer was deposited, followed by a 500 nm gold layer (Discovery 18 Sputtering System, Denton Vacuum LLC, Moorestown, NJ, USA). The tape masks were then removed, leaving a transparent circular window. Devices were diced to size using an automated dicing saw (DISCO 3220).

The LiNbO_3_ transducer was operated in thickness mode, with a fundamental resonance frequency of 6.776 MHz measured by laser Doppler vibrometry (UHF-120SV, Polytec, Waldbronn, Germany). Stimulus frequency, duration, and acoustic pressure were controlled using a signal generator (WF1967, 200 MHz single-channel multifunction generator, NF Corporation, Yokohama, Japan) and a power amplifier (403LA, Electronics & Innovation, Ltd., Rochester, NY, USA). For *in vivo* mouse ECG experiments, a lead zirconate titanate (PZT) ceramic disc transducer (16.7 mm diameter, 3 mm thickness; 840 material, Navy I; American Piezo Ceramics International Ltd.; #70–1000, 2242-D) with a resonant frequency of 668 kHz was used. Stimulus frequency, duration, and acoustic pressure were controlled using a waveform generator (Keysight Technologies, 33500B series) and a power amplifier (TC Power Conversion, AG series amplifier).

### Quantification and statistical analysis

Statistical analyses were performed using GraphPad Prism and MATLAB. All statistical tests were two-tailed. For comparisons between two independent groups, either a Mann–Whitney test or an unpaired *t* -test was used, as indicated. For comparisons among multiple groups, analyses were performed using one-way analysis of variance (ANOVA) followed by Tukey’s multiple-comparisons test, or using a Wilcoxon matched-pairs signed-rank test followed by a Holm–Šidák correction, as indicated. Statistical analyses specific to calcium imaging are described in the corresponding Methods section. No statistical methods were used to predetermine sample sizes for individual experiments. Unless otherwise noted, all experiments and micrographs were repeated at least three times using independent biological replicates.

## DATA AVAILABILITY

All data supporting the findings of this study are available within the paper and its associated files. Source data are provided with this manuscript. Publicly available sequence data (NCBI RefSeq ID: XP_016869435.1) were used in this study.

## ACKNOWLEDGMENTS

The work presented here was generously supported by a research grant from the W.M. Keck Foundation and by the Office of Naval Research (via Grant 13423461) to J. Friend. S. Chalasani is grateful for the support from Mathers Foundation via grant MF-2112-02316 in support of this work. F. Sheikh acknowledges the support of the National Institutes of Health via grants HL162369 and HL181001 for this work. The University of California San Diego Neuroscience Microscopy Shared Facility used in this study is supported by a grant from the NIH (P30 NS047101). The authors are grateful to the University of California and the NANO3 facility at UC San Diego for provision of funds and facilities in support of this work. This work was performed in part at the San Diego Nanotechnology Infrastructure (SDNI) of UCSD, a member of the National Nanotechnology Coordinated Infrastructure, which is supported by the National Science Foundation (Grant ECCS–1542148). The authors are grateful to Megan Anderson, Yanlin Liu, and Wen Mai Wong for their assistance and advice.

