## Supplementary Figures S1 and S2 for "Sonogenetic control of cardiomyocytes and cardiac pacing using exogenous Transient Receptor Potential A1 channels"

**SUPPLEMENTARY INFORMATION**

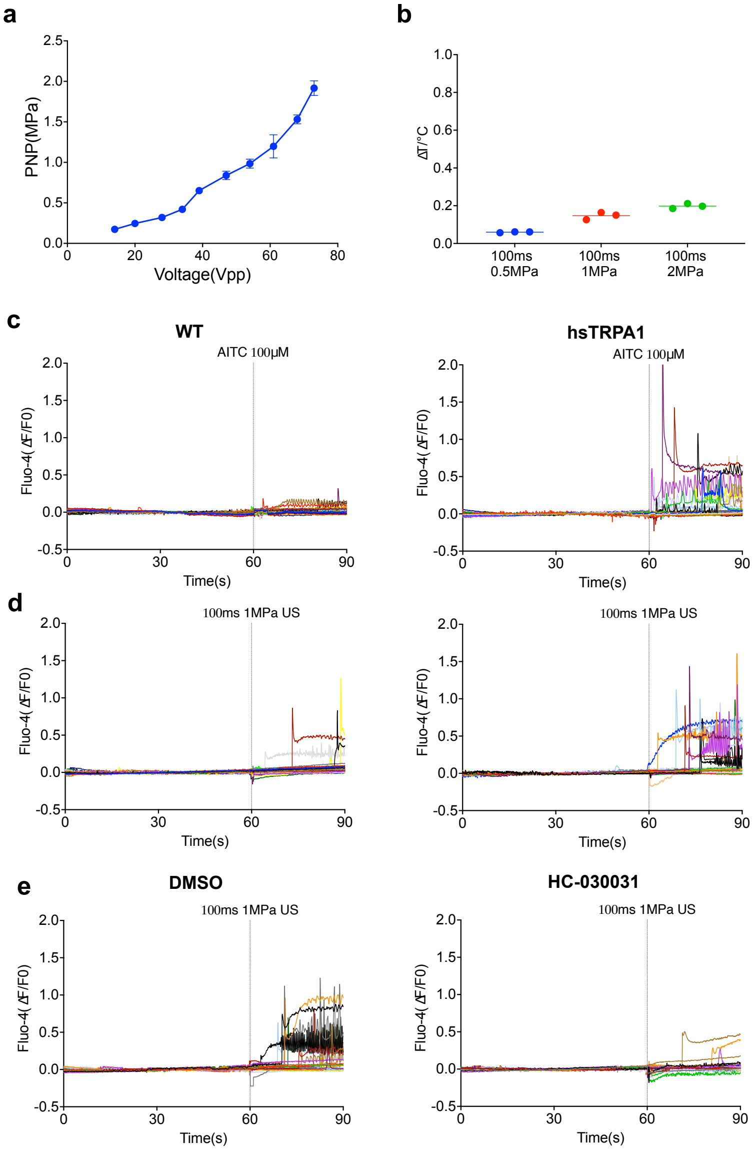


FIG. S1. a, Pressure profile of the ultrasound transducer used for in vitro experiments. Peak negative pressure was measured at a consistent location relative to the face of the transducer through water. Transducer pressure output increased as a function of changing the input voltage. b, Plot showing maximum temperature increases under different ultrasound stimulation parameters. n = 3 assays/condition. c, Fluo-4 fluorescence change trace over time for isolated ventricular cardiomyocytes from WT mice and AAV9-hsTRPA1 mice with treatment of AITC. d, Fluo-4 fluorescence change trace over time for isolated ventricular cardiomyocytes from WT mice and AAV9-hsTRPA1 mice with 100ms 1MPa US stimulation. e, Fluo-4 fluorescence change trace over time for isolated ventricular cardiomyocytes from WT mice and AAV9-hsTRPA1 mice with treatment of DMSO or HC-030031 under 100ms 1MPa US stimulation. Each trace represents one single cell(c,d,e).

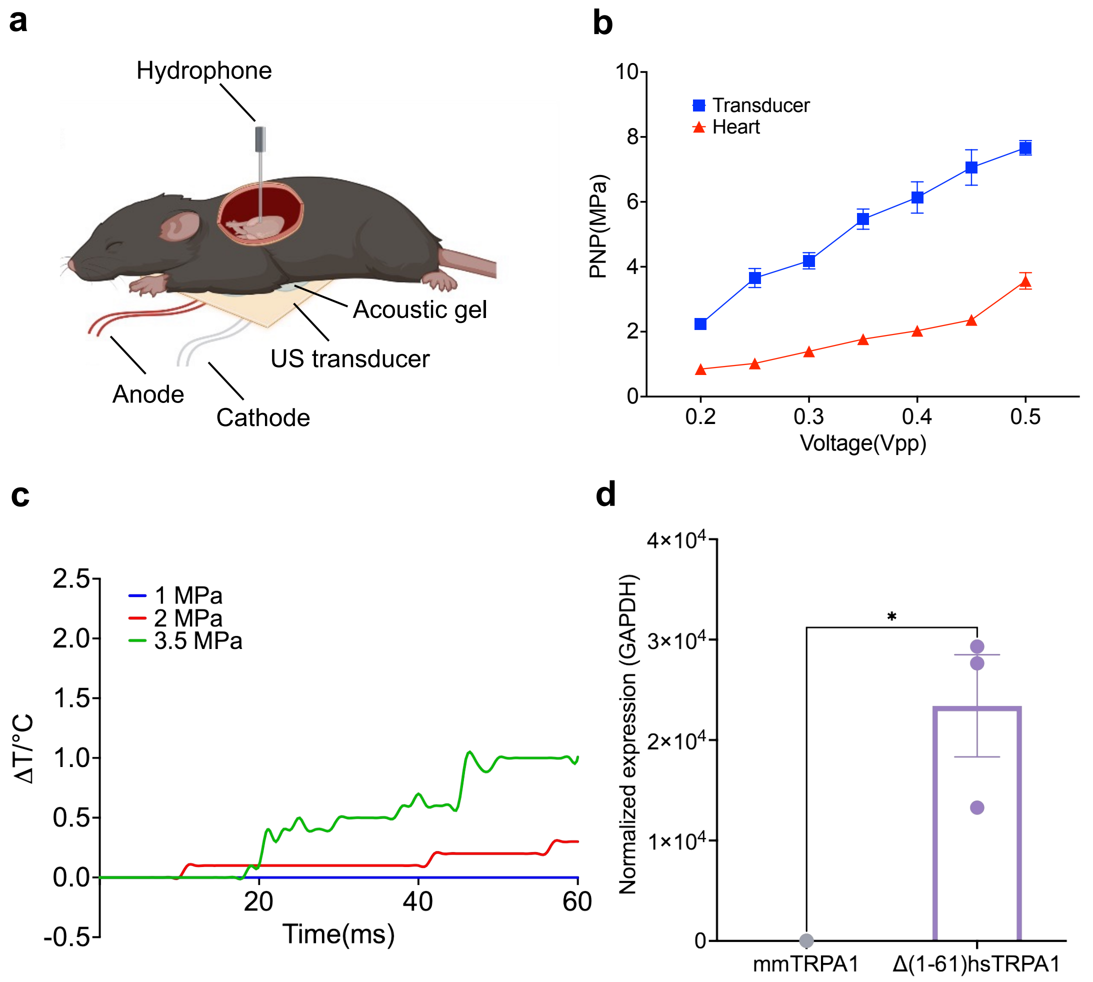


FIG. S2. a, Schemata of hydrophone measurement of in-vivo set-up. b, Pressure measurement of bare transducer and through mice chest with different power input. c. The temperature change measured at the heart front position for in-vivo set-up over time with different ultrasound pressure. Ultrasound pulse is 100ms per 5s. d, RT-qPCR analysis of mmTRPA1 and Δ1-61 hsTRPA1 levels in AAV9-Δ1-61 hsTRPA1 injected wild-type mice. Results are normalized to GAPDH. Data are presented as mean±s.e.m. two-tailed unpaired t test was performed. $*P < 0.05$.
